# Consistency in predicting functions from anatomical and functional connectivity profiles across the cortical cortex

**DOI:** 10.1101/247130

**Authors:** Dongya Wu, Lingzhong Fan, Tianzi Jiang

## Abstract

More and more studies had used connectivity profiles to predict functions of the brain. However, whether anatomical connectivity can predict functions consistently with functional connectivity in various functional domains and whether the connectivity-function relationship is universal across the whole cortex are unknown. Using a linear model, we discovered that anatomical connectivity was comparative to functional connectivity in explaining the variance of functions in most cortical regions, with the exception that anatomical connectivity had poor explaining abilities in brain areas which had high individual task variations. In addition, anatomical connectivity were not that good at capturing individual functional differences and had less inter-subject variation than functional connectivity, however anatomical connectivity could be regarded as more stable in the perspective of parcellation. The current results provided the first comprehensive picture of the relationships between functions and connectivity in the whole human cortex at a fine-grained brain atlas.

## Introduction

Functions of the brain are thought to be constrained by its connectivity profile and one of the major challenges in neuroscience is to establish comprehensive and quantitative connectivity-function relationship across the brain. One critical connectivity profile of a brain region is the extrinsic anatomical connection, known as the region’s connectional fingerprint^1^, and is regarded to be the basis of functional localization in the cortex. Another critical connectivity profile is the functional connectivity at rest and had been corroborated to have the same patterns of functions at task^2, 3^.

The connectivity-function relationship had been studied before and there had already been evidences for a close relationship between connectivity and function. In fact, the boundaries of distinct regions characterized by anatomical connectivity profiles had been found to coincide with boundaries of functionally distinct regions^4, 5, 6, 7^ and anatomical connectivity characterized by fiber bundles had been used to predict functional cortical regions of interests^8^, corroborating that brain regions with different connectivity features should also be functionally distinct. In addition, it had also been further suggested that variations in the connectivity profile should explain functional differentiation^1, 9^ and connectivity features could be used to predict functions. On the one hand, Saygin et al.^10^ used anatomical connectivity determined via diffusion tensor imaging (DTI) to predict functional activation to faces in the fusiform gyrus. Then, Osher et al.^11^ extended it to multiple visual categories across the whole cortex. The close relationship between connectivity and function was demonstrated in the study of Saygin et al.^12^, which showed that anatomical connectivity arose early in development and instructed subsequent functional development of the visual word form area. On the other hand, Tavor et al.^13^ pointed out that it was possible to use functional connectivity at rest to predict individual differences in functional response during tasks.

However, the investigation of the relationship between functions and anatomical connectivity had only been limited to a few visual contrasts in Saygin et al.^10^ and Osher et al.^11^, and it is unknown whether both anatomical and functional connectivity profiles have consistent relationships with functions in various functional domains. Moreover, it remains to address whether the relationship between anatomical connectivity and functions is universal across the whole cortex, especially some high-level regions that are highly variable across individuals.

The current study aims at addressing the above two questions by using a linear regression model and the Brainnetome Atlas^14^ to investigate the connectivity-function relationship at a voxel-wise scale in each cortical region of the Brainnetome atlas across seven tasks from the Human Connectome Project (HCP). Specifically, the linear model was formalized as: Y = Xβ + E, where Y represented the functional activation of voxels in a region, X represented voxels’ connectivity profile and E represented the functional portion that could not be explained by connectivity profile. Anatomical connectivity was characterized by probabilistic tractography from DTI data and functional connectivity was characterized by the correlation of two resting state functional magnetic resonance imaging (fMRI) time series. A portion of E can be controlled by the functional activation level since high-activation voxels have more meaningful information compared to random noises. To prevent over fitting, we divided the subjects into two groups, the training group was used to learn the relationship between the function and connectivity, and the testing group was used to assess the generalization of the learned relationship by evaluating the similarity between the predicted activation and the actual activation. We did permutation test on the linear model by shuffling the parings between Y and X to generate random models. We assessed the individual task variation of each region in a task by averaging the similarities between all pairs of subjects and then subtracted from one.

## Results

### Original model against random models

If a subject’s predicted accuracy from the original model was greater than the 95th percentile of that subject’s random predictions, we regarded it as a better predictor than random. For each contrast and each region, we calculated a permutation indicator as the percentage of test subjects that had connectivity models that performed better than random. For each contrast, we plotted the permutation indicator for each region against that region’s averaged prediction accuracy. We selected some representative contrasts in Fig. S1A for the functional connectivity and Fig. S1B to show the anatomical connectivity. We found that the permutation indicator was highly correlated with the prediction accuracy. Not all the regions’ prediction accuracies were significantly better than the random results; only models that had high accuracies were likely to be better than the random models. Which regions had models with a high accuracy depended on the contrast. Of all the cortical regions across all the test subjects, only a few contrasts’ predictions, such as the PUNISH-REWARD contrast for GAMBLING, were not better than random because these contrasts had a very low activation level throughout the whole cortex.

### Prediction results visualized on the cortex

We selected two representative contrasts and visualized the actual activation and predicted activation of a single subject in Fig. 1 and Figs. S2, the overall patterns of both predicted activation were similar to the actual activation. To analyze the prediction results quantitatively, we presented the each region’s activation profile and individual task variation in Fig. 2 and Fig. S3. The activation profile of a region was characterized as the averaged activation of all the voxels within that region across all the training subjects. The lateral pre-frontal regions such as the inferior frontal junction (IFJ) and inferior frontal sulcus (IFS), and a few visual cortices such as the caudal lingual gyrus (cLinG) and middle occipital gyrus (mOccG) showed high individual task variations across all tasks. We also presented the averaged prediction accuracies made by both connectivity profiles in Fig. 3 and Fig. S4. Overall, the patterns of both prediction accuracies were very similar and both connectivity profiles had good prediction results in most activated regions shown in Fig. 2. However, anatomical connectivity had poor prediction results compared to functional connectivity in some pre-frontal and visual cortices which had relatively high individual task variation. The precise descriptions of these results were presented below.

**Figure 1.**
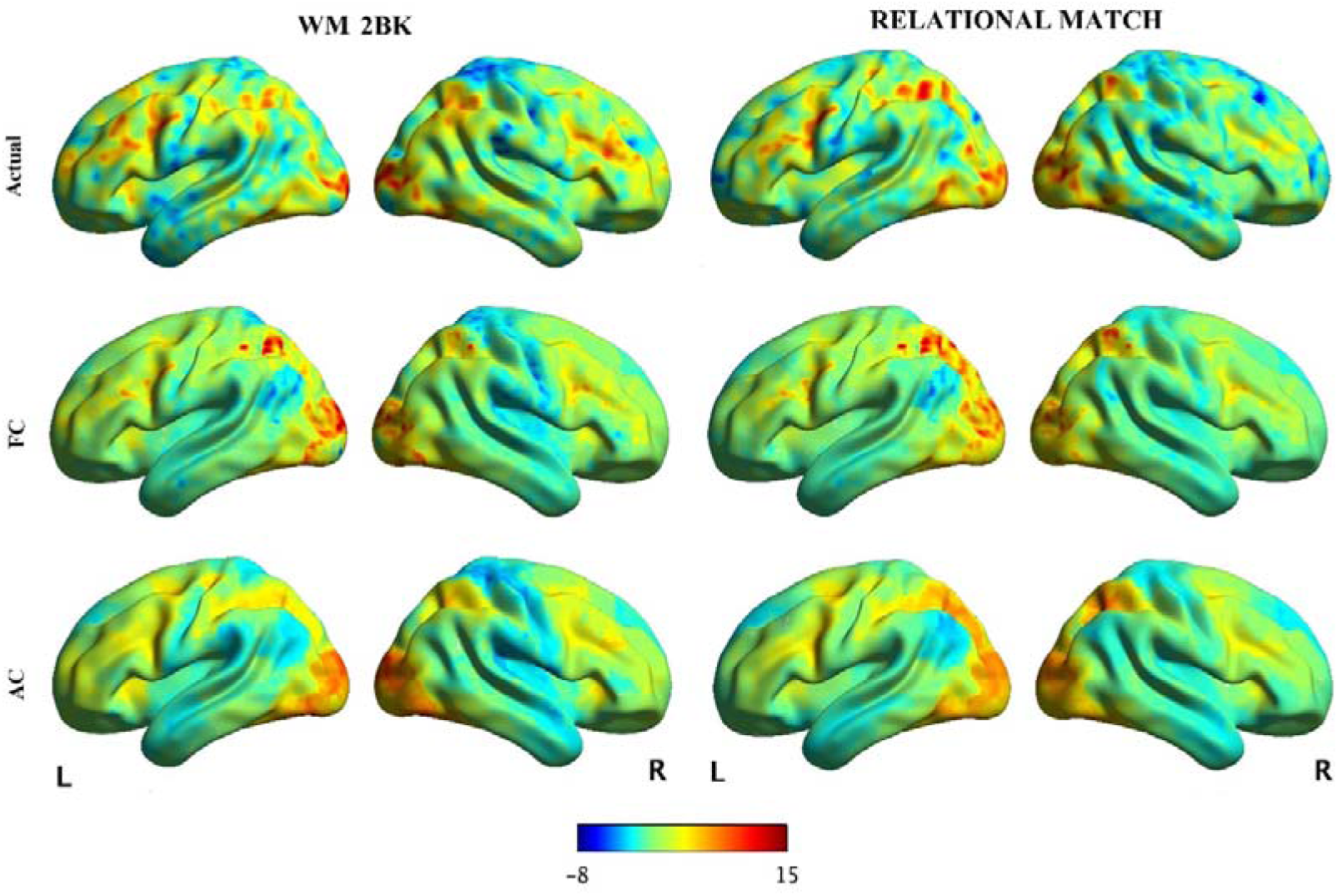
A single subject’s prediction result visualized on the cortex. We selected two contrasts and visualized the actual and predicted activation of a single subject on the cortex. More examples were shown in Fig. S2. AC represents the activation made by anatomical connectivity and FC represents the activation made by functional connectivity. The overall patterns of predicted activation by both connectivity profiles were very similar to the actual activation. The quantitative analyses of the prediction results were presented in the following.

**Figure 2.**
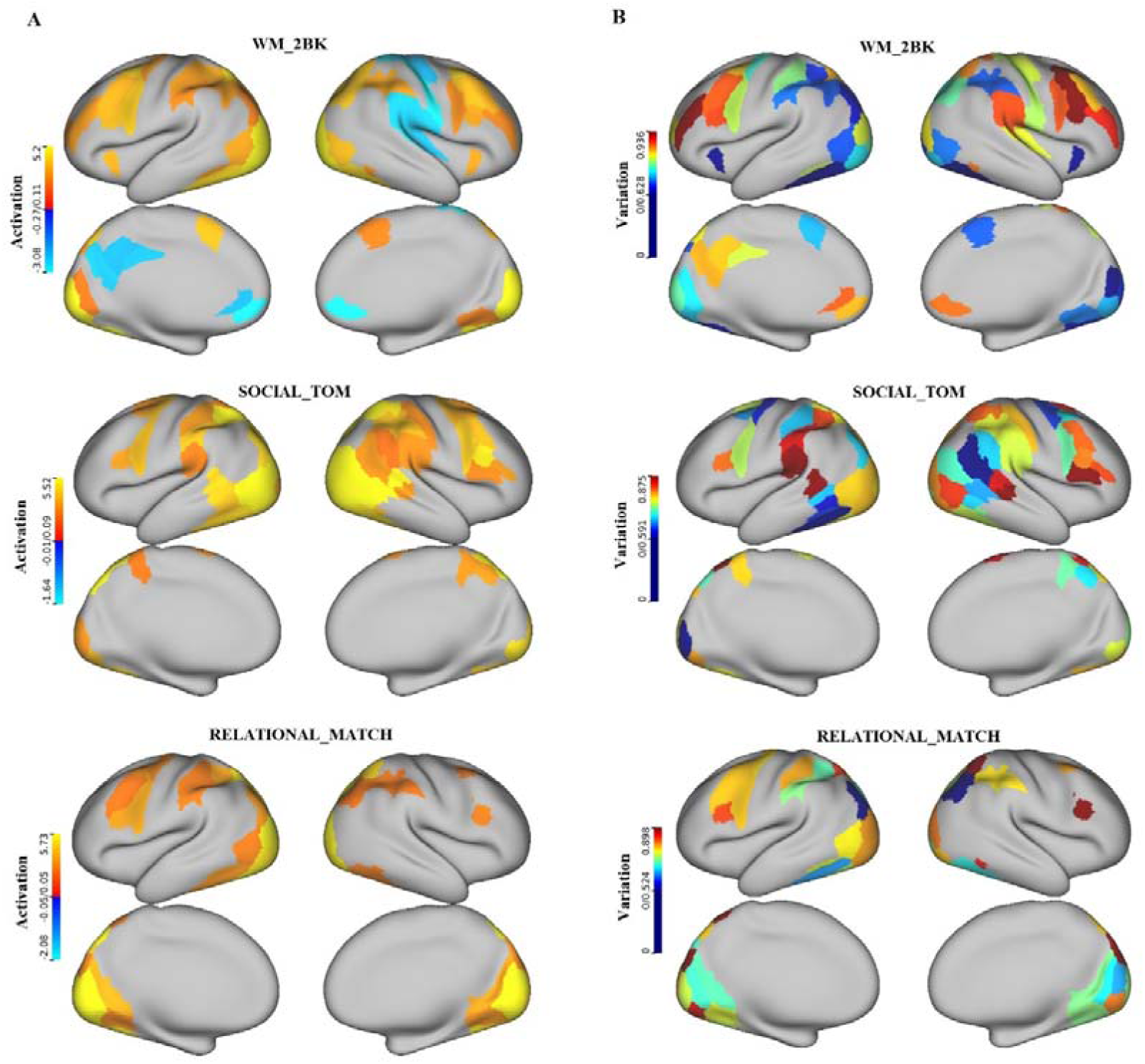
Activation profile and individual task variation. (A) Each region’s mean activation profile were calculated by first averaging each voxel’s activation level within a subject and then averaging each subject’s mean activation. The threshold values were determined by one sample t-test to find regions that had mean activation significantly different from zero (*p* < 0.05) across subjects. (B) The individual task variation of region’s that had mean activation level above the threshold were shown. The pre-frontal regions such as the IFJ and IFS, and a few visual cortices such as the cLinG and mOccG showed high individual task variations across all tasks.

**Figure 3.**
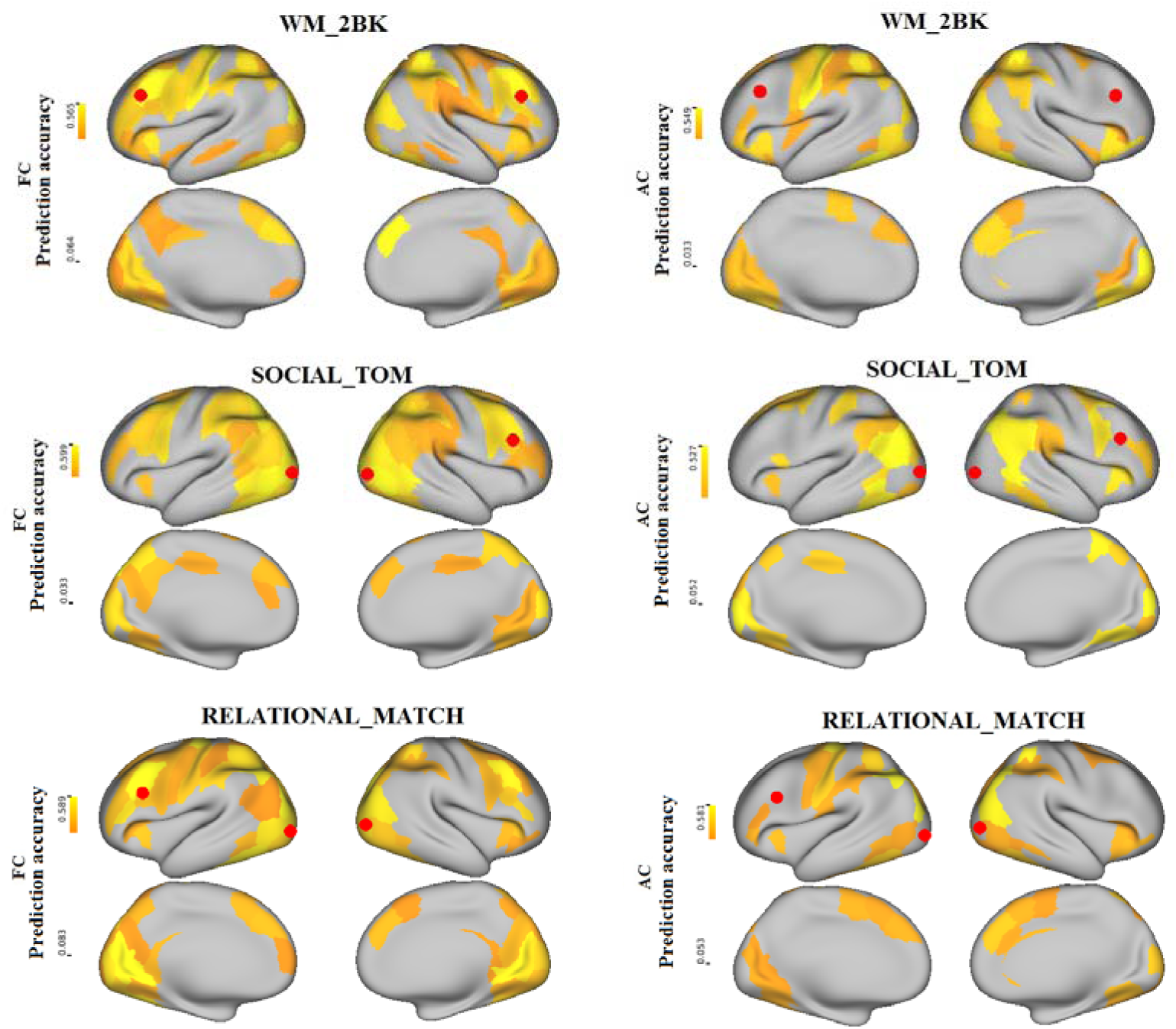
Averaged prediction accuracies visualized on the cortex. We selected some representative contrasts and visualized the the prediction accuracies made by both connectivity profiles on the cortex. Both connectivity profiles had similar prediction accuracies in most regions, however, in a few pre-frontal and visual cortices that had relatively high activation as well as high individual task variation as shown in Fig. 2, anatomical connectivity had poor prediction acuuracies compared to functional connectivity.

### Activation profile and model performance

The model performance of a region was related to that region’s activation profile. We correlated the activation profile of each region with the averaged model prediction accuracy of the same region, and the results were provided in Figs. S5A and S5B. The prediction accuracy was highly correlated with the absolute activation level, indicating that both connectivity profiles could only have good prediction results in activated regions, since random noises had more impacts on the model in less activated regions. However, anatomical connectivity had bad prediction results in a few regions consistently across different tasks, even though these regions had relatively high activation in the tasks. Those regions included a few pre-frontal regions such as the IFJ and IFS, and a few visual cortices such as the cLinG and mOccG.

### Model performance comparison between anatomical and functional connectivity

For each contrast, we plotted each cortical region’s prediction accuracy made using anatomical connectivity against that cortical region’s prediction accuracy made using functional connectivity, as seen in Fig. 4. We did paired-*t* test to find the regions that had big differences of prediction accuracies between these two connectivity profiles. After a strictly family wise error correction with *p* < 1e-4, only a few pre-frontal and visual cortices survived the correction consistently across different tasks. Overall, the prediction accuracies made by anatomical connectivity were very comparative to those made by functional connectivity, indicating that anatomical connectivity had good explaining ability to task activation in most regions across a wide task domains. However, anatomical connectivity had very low prediction accuracies compared to functional connectivity in a few regions such as the IFJ, IFS, cLinG and mOccG across many task domains, even though those regions had relatively high activation. The particular properties of these regions were discussed below.

**Figure 4.**
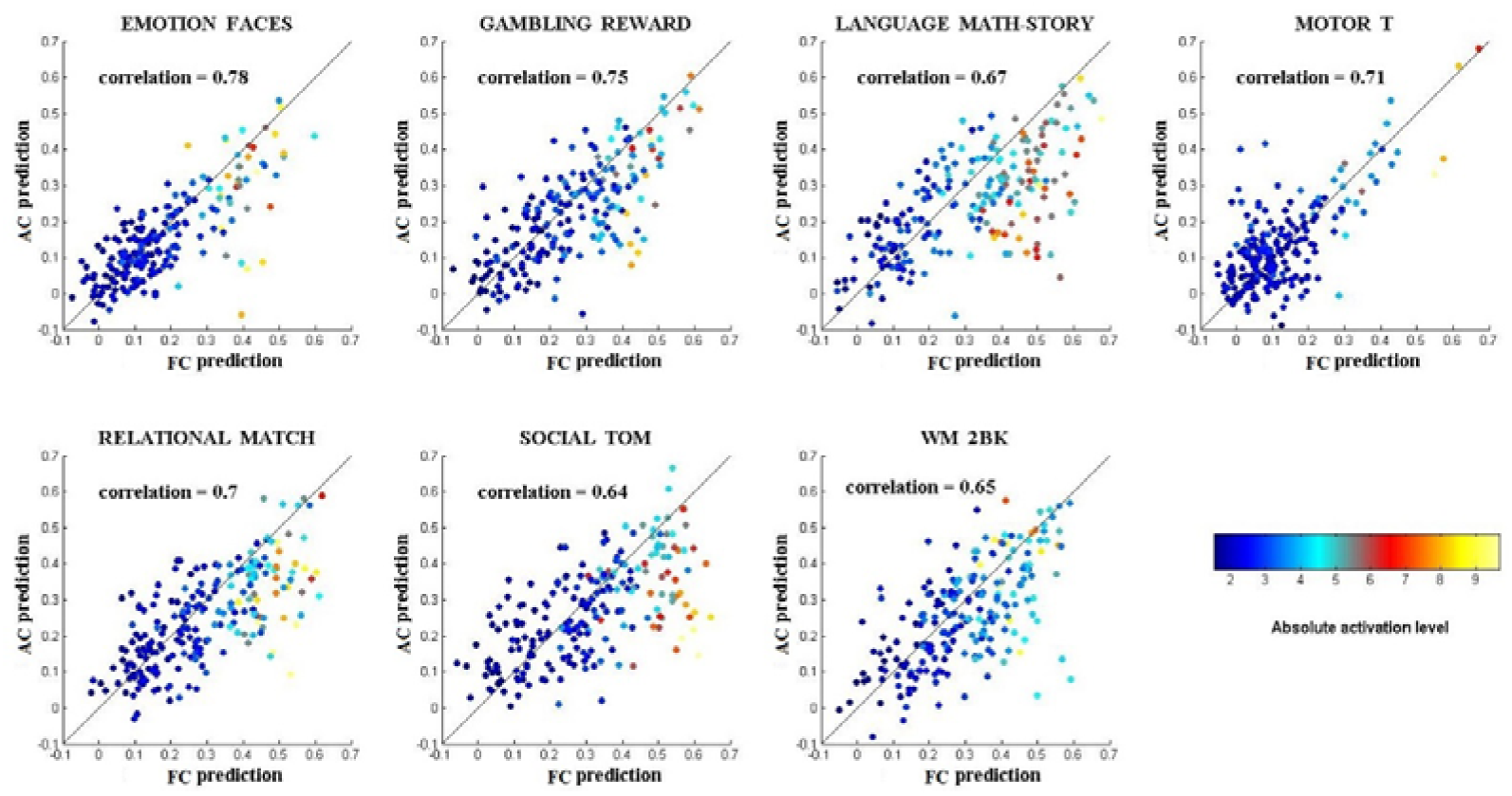
Model performance comparison between two kinds of connectivity features. Each region’s prediction accuracy obtained using anatomical connectivity is plotted against that region’s prediction accuracy obtained by functional connectivity. Each region is also color coded by its absolute activation level. The antidiagonal indicates that the prediction accuracies made by both connectivities are equal. Overall, the prediction accuracies made by these two kinds of connectivity are very similar, but there are also some regions that fall far from the antidiagonal, indicating that the differences between the prediction accuracies of these two connectivity features are great.

### Model performance and individual task variation

We plotted each region’s individual task variation against that region’s prediction accuracy made using anatomical or functional connectivity in Fig. 5 and Figs. S6. Most regions that had more individual task variation were less activated and random noises had more effects in these regions. However, the few pre-frontal and visual cortices mentioned previously had relatively high activation as well as high individual task variation that must be related to other factors beyond random noises. In these regions with relatively high activation as well as high individual task variation, anatomical connectivity had poor explaining ability of the task activation but functional connectivity still made good predictions.

**Figure 5.**
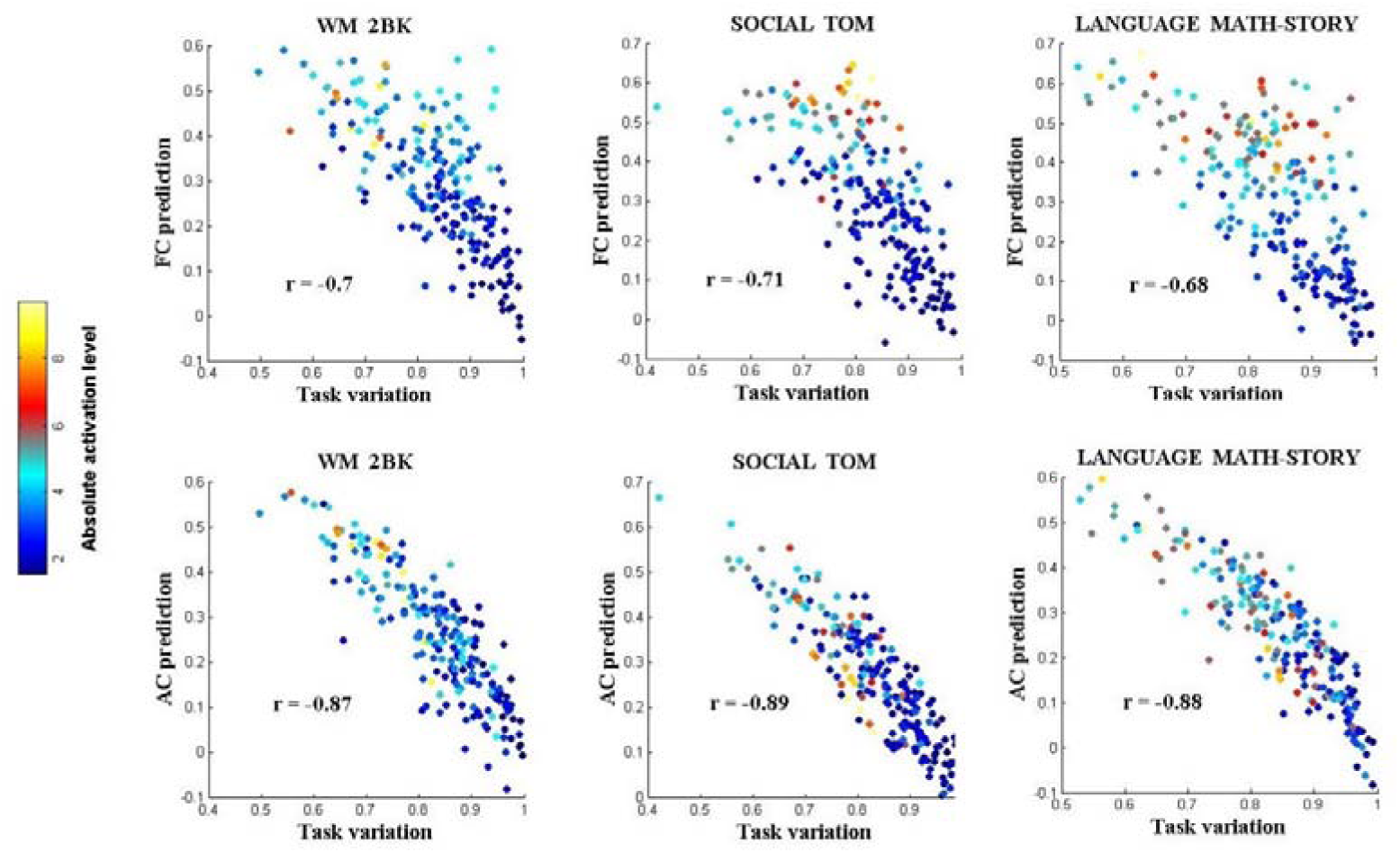
Individual task variation and prediction accuracies. Each region’s individual task variation is highly correlated to that region’s predition accuracy made using connectivity profiles. Most regions that have high individual functional variation also have low activation. A few pre-frontal and visual cortices have high activation as well as high individual functional variation. Anotomical connectivity has poor prediction accuracies compared to functional connectivity in those pre-frontal and visual cortices.

### Individual variation of functions and connectivity features

For each of the 210 cortical regions, we calculated that region’s inter-subject variation for both connectivity features (see Figs. 6A and 6B). We did a paired *t*-test and found that the functional connectivity’s indicator for the inter-subject variation was very significantly smaller (*p* < 1e-6) than that for the anatomical connectivity, a finding which meant that functional connectivity was more variable than the anatomical connectivity across the subjects. The patterns of between-subject variation of functional connectivity but not anatomical connectivity were similar to the patterns of individual task variation in Fig. 2, with motor areas had relatively low between-subject variation and pre-frontal regions had relatively high between-subject variation. We tested how the individual variation of connectivity features were related to the individual variation of functional activation. Since the random noises was a potential confounding factor, we also used the activation level as an explaining factor for the individual variation of function. We concatenated each region’s individual variation of task activation and activation level across all contrasts and regressed out the factor of activation level from individual variation of task activation. We found that the correlation between individual variation of task activation and individual variation of functional connectivity was 0.25 (*p* < 1e-10), but the correlation between individual variation of task activation and individual variation of anatomical connectivity was 0.03 (*p* = 0.005), meaning that the individual variation of task activation could not be related to the individual variation of anatomical connectivity.

**Figure 6.**
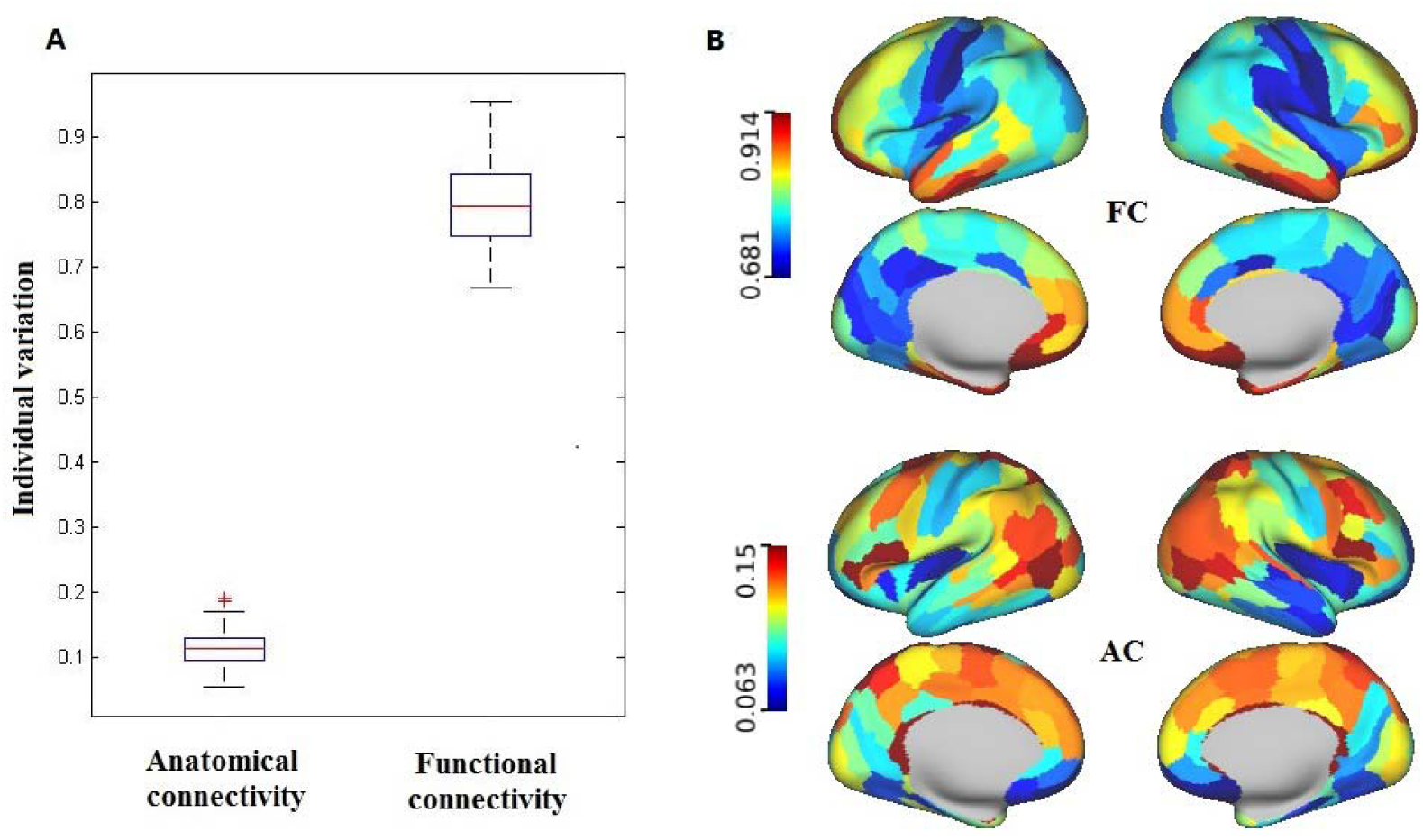
The inter-subject variation of both connectivity features. (A) Each of the 210 cortical region’s indicator of inter-subject variation is plotted in the boxplot. We did a paired *t*-test and found that the functional connectivity’s indicator of inter-subject variation was very significantly smaller (*p* < 1e-6) than that for anatomical connectivity, meaning that functional connectivity was more variable across subjects than anatomical connectivity. (B) Each region’s inter-subject variation of connectivity features is mapped onto the cortex. Functional connectivity has less individual variation in motor areas and more variation in pre-frontal areas. However, the individual variation pattern of anatomical connectivity is not similar to that of functional connectivity.

### Prediction of individual functional difference

We concatenated the prediction for each region to calculate the between-subjects cross-correlation matrices for the whole cortex. We tested whether the between-subjects cross-correlation matrices were diagonal-dominant by calculating the diagonal-dominant indicator as the percentage of test subjects that had predictions that best matched the subject’s own actual response. The between-subjects cross-correlation matrices for the whole cortex for some of the contrasts are presented in Fig. 7. The between-subjects cross-correlation matrices made using functional connectivity are more diagonally dominant than those made using anatomical connectivity, a finding which indicated that anatomical connectivity could not capture as much of the individual functional differences as functional connectivity could and corresponded with the previous result that individual task variations were less correlated with anatomical connectivity profile.

**Figure 7.**
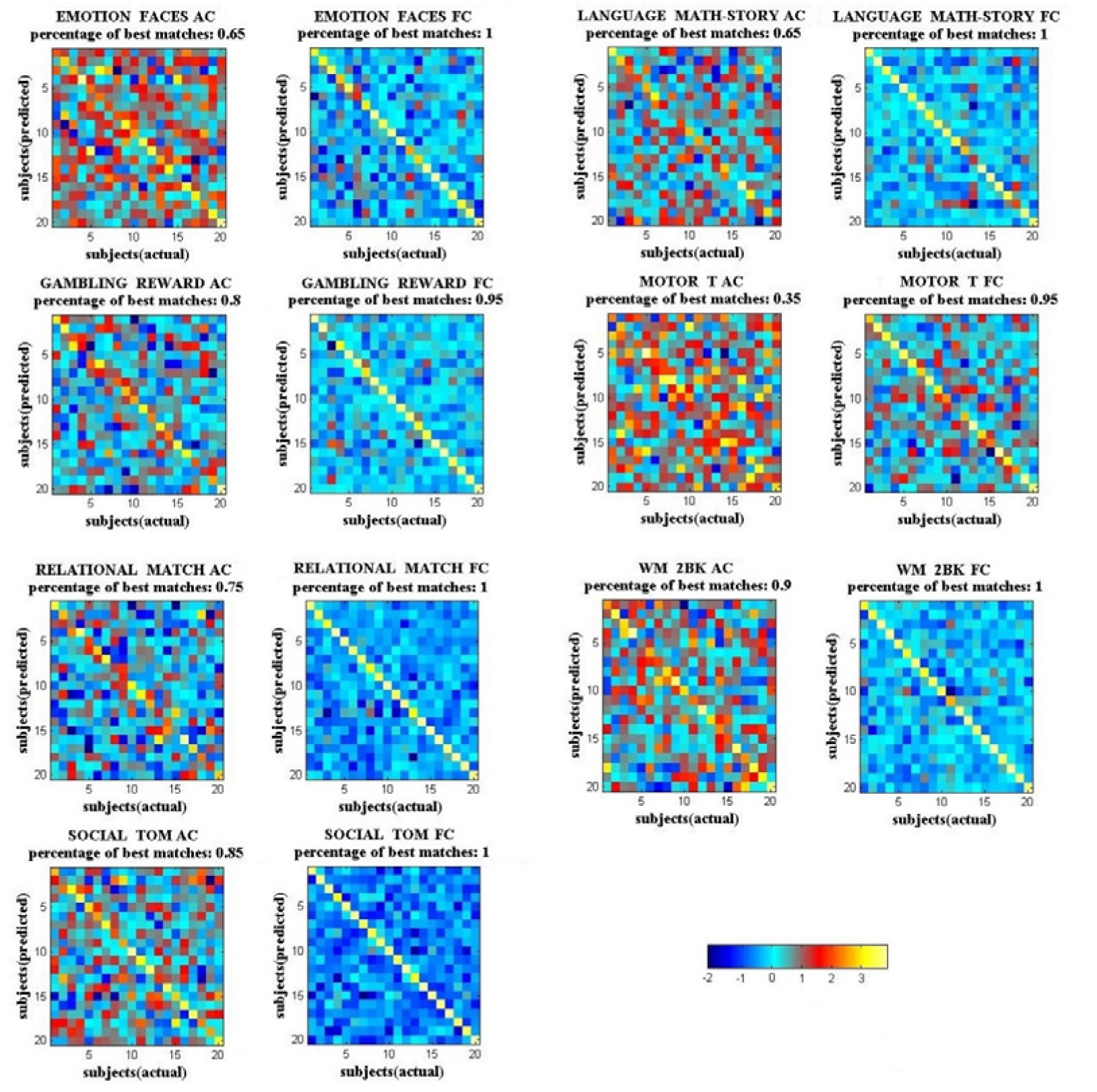
Between-subjects cross-correlation matrices for the whole cortex. The cross-correlation matrices made using functional connectivity are more diagonally dominant than those made using anatomical connectivity. Almost all the subjects’ predictions made using functional connectivity best matched their own actual functional responses, but the predictions made using anatomical connectivity did not capture as many individual differences as functional connectivity.

## Discussion

To investigate the precise relationship between connectivity profile and function, we examined whether both connectivity profiles could be used in a linear regression model to predict functional activation in the whole human cortex across a wide range of task domains. We found that for each contrast, the connectivity profile could only have good predictions of functional activations in cortical regions that were the most task relevant. Task relevant regions are those regions that exhibit high activations during a task. The prediction accuracies of the non-task relevant regions were comparatively much lower and unlikely to be better than random permutation results. Thus, testing the connectivity-function relationship in task-relevant regions is necessary. That is also the reason why we adopted the HCP dataset, which contains a wide range of functional domains. By examining the connectivity-function relationship at a fine-grained brainnetome atlas across seven tasks, we demonstrated that anatomical connectivity had a close relationship with function as well as functional connectivity in most of cortical regions, however in a few regions which had high individual task variation such as the pre-frontal cortices IFJ and IFS, anatomical connectivity made poor predictions compared to functional connectivity. Although anatomical connectivity could have comparative prediction accuracies in most regions, it could not capture individual functional differences as well as functional connectivity.

The results that anatomical connectivity were comparative to functional connectivity in predicting the functions in most cortical regions validated the close relationship between functions of the human brain and its underlying structural substrates. Numerous studies had demonstrated that the wiring of neurons were essential in determining their functions in non-human animals. For example, Briggman et al.^15^ indicated that the structural wiring asymmetry contributes to the direction selectivity of the retina. And Glickfeld et al.^16^ also showed that cortical-cortical projections were critical in the transmission of specific information to downstream targets, demonstrated that specific connectivity patterns were important for areal specialization, where neighboring neurons were functional diverse. Therefore, the connections of neurons or regions determined their information input and output flow, and were critical for defining their functional properties, which were the fundamental underlying reasons that we found a close relationship between connectivity and functions in the human brain at the neuroimaging level.

A plausible explanation of why functional connectivity could reflect individual functional differences better than anatomical connectivity is the close relationship of functional networks during rest and tasks. As others have showed, brain regions that interact with each other during tasks are also continuously interacting with each other during rest with the same functional hierarchy, and the functional network architecture during tasks is shaped primarily by an intrinsic network architecture that is also present during rest^2, 3^. Mennes et al.^17^ found out that a region’s intrinsic activity as measured during rest predicted that same region’s activity induced by a Flanker task and Tavor et al.^13^ also pointed out that resting state connectivity profile can be used to capture the individual differences of task activation in a wide task domains.

Of course, functional networks reflected the underlying structural networks^18^. Measurements of spontaneous activity revealed functional connectivity patterns that were similar to anatomical connectivity, suggesting that functional connectivity was highly constrained by anatomical connectivity^19, 20^. Simulation studies had also been done to predict human resting-state functional connectivity from anatomical connectivity^21^. However, although structural networks underlie functional networks, the relationship between structural and functional connections is far more complex. As Honey et al.^21^ pointed out, strong functional connections commonly existed between regions with no direct structural connection, and indirect connections and interregional distance accounted for some of the variability in functional connectivity that was unexplained by direct anatomical connectivity. Therefore, functional connections reflect a combination of numerous dynamic influences traveling through the network along many paths, some of which are indirect and take multiple intermediate steps^22^. In this view, using only the structural connections of a cortical region may not capture these indirect and dynamic influences as well as functional connections do.

In addition, the results that anatomical connectivity was less variable across subjects than functional connectivity and reflected less individual functional differences could also be interpreted from another perspective. Individual variations are seen in all task domains, however, the individual differences in task activations are often related to volatile factors and are removed through averaging. Therefore, even though anatomical connectivity could not reflect individual functional differences as well as functional connectivity due to the close relationship between the functional networks during tasks and at rest, anatomical connectivity could be regarded as more stable than functional connectivity and would have advantages in other perspectives, such as parcellation of the brain, which demand the borders of the parcellation to be smooth and less variable.

Further, there were also some limitations in characterizing anatomical connectivity and functions in this research. The anatomical connectivity derived from diffusion MRI tractography in this research. However, Thomas et al.^23^ compared tract-tracer and diffusion MRI tractography and observed limited accuracy of tractography, and Reveley et al.^24^ further validated that tractography methods were biased in their identification of long-range projections toward cortical gyri, much of which could be ascribed to the superficial white matter. Maier-Hein et al.^25^ also showed that the reliability of fiber tracking algorithm was limited based on orientation information alone and this need to be considered when interpreting tractography and connectivity results. Besides, we used connection propabilities to characterize anatomical connectivity strength and the accurate defining of the connectivity strength also remained open. The functions at task state and functional connectivity at resting state were both characterized by blood oxygen-level dependent (BOLD) signal. The BOLD signal reflected the underlying neural connectivity indirectly by assessing the changes in deoxyhemoglobin, and its signal source can be complex and is also dependent on imaging parameters and techniques^26^. The extent to which that BOLD signal could reflect the actual underlying neuronal functions was also obscured^27^. Therefore, these limitations must be considered when interpreting the results.

Even though resting state functional connectivity and functional activation both derived from BOLD signal, we demonstrated that extrinsic anatomical connections could as well consistently predicted the functions of most cortical regions across a wide range of behavioral domains. The inconsistency exhibited in a few pre-frontal and visual cortices with high individual variations and these high individual variations could not be explained by anatomical connectivity. Future works could also explore the origins of these high inter-subject variations and build more complex models which incorporate indirect anatomical connections to reflect the dynamic influences of functional networks.

## Methods

### Human Connectome Project data

We used the minimally pre-processed data^28^ provided by the HCP. In total, a group of 40 subjects were used in our analyses. Acquisition parameters and processing are described in detail in several publications^29, 30, 31, 32^.

Briefly, DTI data were acquired using single-shot 2D spin-echo multiband echo planar imaging on a Siemens 3 Tesla Skyra system^31^. These consist of 3 shells (b-values=1000, 2000, and 3000 s/mm2) with 270 diffusion directions isotropically distributed among the shells, and six b=0 acquisitions within each shell, with a spatial resolution of 1.25 mm isotropic voxels. Each subject’s diffusion data had already been registered to his or her own native structural space^28^.

Resting and task fMRI scans were acquired at 2 mm isotropic resolution, with a fast TR sampling rate at 0.72 s using multiband pulse sequences^32^. Both sets of functional data had already been registered to MNI space^28^. Each subject had four 15-minute resting fMRI runs, with a total of 1,200 time points per run. The resting fMRI data were further pre-processed by FIX to automatically remove the effect of structured artefacts^33, 34^. The task fMRI contained 86 contrasts from seven task domains, labeled as EMOTION, GAMBLING, LANGUAGE, MOTOR, RELATIONAL, SOCIAL, and WM (working memory). The details of the tasks are described in Barch et al.^29^. Because most of the contrast maps were paired with a related negative contrast, which is redundant for the purpose of regression modeling, we excluded those redundant contrasts and kept 47 contrasts for further regression analysis.

### Connectivity profile

We calculated two kinds of connectivity profiles using a brain atlas^14^ containing 210 cortical regions and 36 sub-cortical regions in our analyses. We tested the connectivity-function relationship on each of the 210 cortical regions individually using regression analysis. Each of the 210 cortical regions was used as a seed region. For every seed region, therefore, there were 209 target cortical regions and 36 target sub-cortical regions. All the voxels within the seed region were characterized by two kinds of connectivity features of 245 dimensions, representing the connectivity of each voxel in the seed region to the remaining 245 target regions. Even though we tested the connectivity-function relationship only on the cortical regions, the connectivity profiles also included sub-cortical features.

The functional connectivity was calculated from resting fMRI data. The four runs of individual resting state time series data were concatenated after being demeaned and variance normalized. The voxel time series for every voxel in the seed region was correlated with the averaged time series for each of the remaining 245 target regions.

The anatomical connectivity was determined via probabilistic diffusion tractography. Fiber orientations were estimated per voxel, and probabilistic diffusion tractography was performed using FSL-FDT^35^ with 5000 streamline samples in each seed voxel to create a connectivity distribution to each of the remaining 245 target regions, while avoiding a mask consisting of the ventricles. The probabilistic tractography was performed on each subject’s native space, but the atlas and functional data were in MNI space. The nonlinear volume registration between MNI and native space is described in Glasser et al.^28^.

### Model training

To avoid over fitting, we separated the 40 subjects into two groups of 20 subjects. One group was used to train the regression model; the other was used to test the regression model and assess the connectivity-function relationship. For each of the total 47 contrasts, we performed a regression analysis on each of the 210 cortical regions using both connectivities. The regression analysis was modeled as: Y = Xβ + E, where Y is the *t*-statistical value of the contrast maps; X represents either functional connectivity or anatomical connectivity; β is the regression coefficient to be estimated from the regression model. To train the regression model on the ith cortical region, we concatenated all the voxels of each training subject’s ith cortical region into a column. Assuming the ith cortical region has *n_i_* voxels, Y is a single column vector of length *N_i_* = 20 × *n_i_* representing the functional response of all the seed voxels, X is a matrix of *N_i_* columns and 246 rows representing the connectivity features of all the seed voxels plus an intercept term, β is a single column vector of length 246 representing how each connectivity feature contributes to predicting a seed region’s functional response. Similarly, we also obtained a connectivity matrix of *N_i_* columns and 246 rows from the testing group. After estimating the regression model’s coefficients from the training group, we applied these coefficients to the testing group’s connectivity matrix to get the predictive functional response of the subjects in the testing group. To get the predictive functional response of the entire cortex, we repeated the same procedure for every cortical region and then concatenated every region’s prediction.

### Model assessments

We correlated the predicted response of every subject in the testing group with the actual functional response of the same subject to evaluate the accuracy of the predictions. We used the correlation coefficients (*r*) to assess the prediction instead of the mean squared error (MSE) or mean absolute error (MAE) because, unlike MSE and MAE, which are not standardized and un-bounded, *r* is standardized and bounded between 0 and 1. In addition, under the least square conditions of the regression model, *r* squared equals *R*^2^, which is the goodness of fit of the model and represents the proportion of the variance in the fMRI response that can be explained by the connectivity features.

To evaluate whether the model could capture individual fMRI response differences, we correlated each testing subject’s predicted response with each testing subject’s actual response to form between-subjects cross-correlation matrices. Theses matrices are row- and column-normalized, which accounts for the fact that the actual responses are more variable than the predicted responses and removes the global mean correlation, as stated in Tavor et al.^13^. If the matrix is diagonally dominant, which indicates that each subject’s predictive response best matches that subject’s actual response, then the connectivity profile can capture individual specificities in fMRI activations.

We also compared the inter-subject variation of these two connectivity features for each of the 210 cortical regions. For each region, we transformed each subject’s connectivity matrix into a column vector and concatenated all the training subjects’ connectivity features for that region across the column to get the across-subjects’ connectivity matrix. We calculated the indicator of the inter-subject variation as the percentage of the greatest singular value of the across-subjects’ connectivity matrix against the sum of all the singular values of the across-subjects’ connectivity matrix. If the connectivity feature has low inter-subject variation, the across-subjects’ connectivity matrix should be highly degenerate and the indicator will be close to one. If the connectivity feature has high inter-subject variation, the across-subjects’ connectivity matrix should be less degenerate and the indicator will be close to zero.

### Permutation test

Also, we did 1000 random permutations to test the connectivity model. We trained the models in the same manner, but the pairings between each voxel’s connectivity feature and its functional response were shuffled. We then tested how these random models performed on the testing group. To test whether the performance of the original model was statistically meaningful, we compared the performance of the original model with the performance of the random models. For each test subject, we got one prediction accuracy from the original model and 1000 prediction accuracies from the random models and calculated whether each subject’s prediction accuracy from the original model was higher than the 95th percentile of that subject’s random predictions. One thing to notice here is that we only shuffled the data in the training group, but not in the testing group. Since we trained the regression model one region at a time, we shuffled the pairings within the seed region, but not across the whole cortex.

## Authors’ contributions

Jiang and Wu designed the research, Wu performed the analysis, Jiang, Wu and Fan drafted the manuscript.

## Acknowledgement

We appreciate the editing assistance of Rhoda E. and Edmund F. Perozzi, PhDs

